# TrueProbes: Quantitative Single-Molecule RNA-FISH Probe Design Improves RNA Detection

**DOI:** 10.1101/2025.08.14.670355

**Authors:** Jason J. Hughes, Benjamin K. Kesler, John E. Adams, Blythe G. Hospelhorn, Gregor Neuert

**Affiliations:** Department of Molecular Physiology and Biophysics, School of Medicine, Vanderbilt University, Nashville, TN 37232, USA; Vanderbilt Genetics Institute, School of Medicine, Vanderbilt University, Nashville, TN 37232, USA; Department of Biomedical Engineering, School of Engineering, Vanderbilt University, Nashville, TN 37232, USA; Department of Pharmacology, School of Medicine, Vanderbilt University, Nashville, TN 37232, USA; Center for Computational Systems Biology, School of Medicine, Vanderbilt University, Nashville, TN 37232, USA

## Abstract

Single-molecule RNA fluorescence in situ hybridization (smRNA-FISH) is a widely used method for visualizing and quantifying RNA molecules in cells and tissues at high spatial resolution. The technique relies on fluorescently labeled oligonucleotide probes that hybridize to target RNA. Accurate quantification depends on high probe specificity to ensure fluorescent signals reflect target RNA binding rather than off-target interactions. Numerous factors, including genome sequence complexity, secondary probe structure, hybridization conditions, and gene expression variability across cell types and lines, influence smRNA-FISH probe efficacy. Existing smRNA-FISH probe design tools have limitations, including narrow heuristics, incomplete off-target assessment, and reliance on “trial-and-error approaches. To address these challenges, we developed TrueProbes, a probe design software platform that integrates genome-wide BLAST-based binding analysis with thermodynamic modeling to generate high-specificity probe sets. TrueProbes ranks and selects probes based on predicted binding affinity, target specificity, and structural constraints. It also incorporates a thermodynamic-kinetic simulation model to provide predictive design metrics and optimize probe performance under user-defined conditions. We benchmarked TrueProbes against several widely used smRNA-FISH design tools and found that it consistently outperformed alternatives across multiple computational metrics and experimental validation assays. Probes designed with TrueProbes showed enhanced target selectivity and superior experimental performance.

## Introduction

Gene regulation is a fundamental biological process that underpins development and homeostasis, and it is frequently dysregulated in disease (1, 2). Quantification of gene expression has traditionally relied on population-level measurements of RNA or protein, using techniques such as quantitative PCR (qPCR), RNA sequencing, western blotting, and proteomics (3, 4). As a diagnostic tool, these gene expression measurements can distinguish between healthy and diseased cells (2).

For over 25 years, single-cell studies have revealed substantial variability in gene expression, even among cells with identical genotypes (5-8). Recent technological advances now allow for finer resolution of expression differences among cells within the same organ, highlighting the diversity of cell types and transcriptional states (9). Detailed analysis of single-cell data has shown that this variability can be leveraged to infer gene-regulatory mechanisms that are undetectable in bulk measurements (10).

Several approaches can be used to quantify RNA at the single-cell resolution, including single-cell RNA-sequencing, live-cell RNA reporters, and single-molecule RNA fluorescence in situ hybridization (smRNA-FISH). Each has limitations: single-cell RNA-seq suffers from technical noise, low sensitivity, and indirect molecule counting (11), whereas live-cell reporters require the engineering of every transcript of interest (12).

By contrast, smFISH is an imaging method that utilizes multiple oligonucleotide probes either labeled directly with fluorophores or tagged for secondary-probe binding (Figure 1A) to detect individual RNAs (13, 14). Ideally, probes display high homology to targets and minimal complementarity to off-target transcripts that contribute to autofluorescence. smRNA-FISH can measure hundreds to thousands of RNA species with high spatial and temporal resolution in single cells and tissues (15, 16). Accurate quantification requires probe-derived fluorescence to be specific to the intended transcript, as off-target binding can elevate background or create false-positive spots (17).

**Figure 1:**
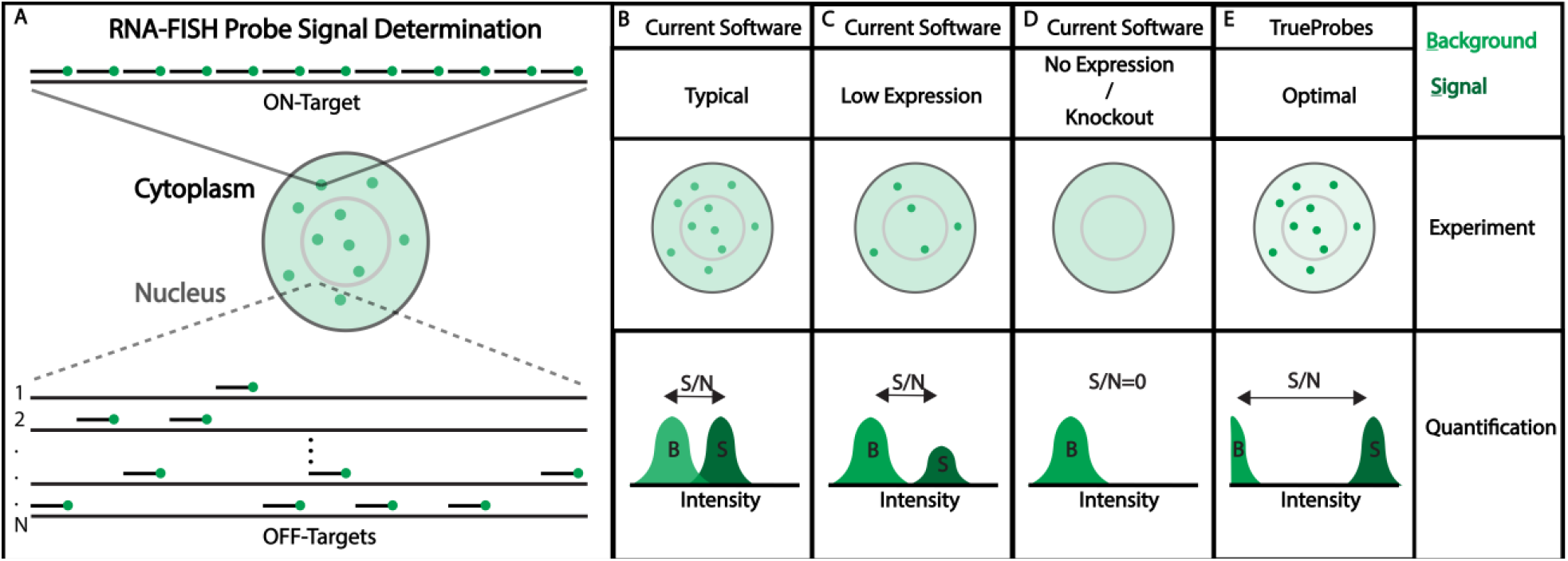
Traditional smRNA-FISH probe design limits sensitivity and specificity. Panels A–E illustrate conceptual smRNA-FISH design scenarios affecting signal-to-noise ratio (SNR). (A) Spot SNR is determined by the number of probes bound to on-target RNA (signal, S) relative to probes bound to off targets and cellular autofluorescence (background, B). On-target and off-target localization are assumed to be random. (B) A typical design shows moderate background and submaximal spot intensity due to suboptimal probe number or affinity, resulting in overlapping signal and background distributions and reduced SNR. (C) In low expression conditions, the background remains constant but fewer on-target transcripts yield fewer detectable spots, reducing overall detection sensitivity. (D) In knockout or no-expression cases, background persists while on-target signal is absent, highlighting the contribution of off-target binding and autofluorescence. (E) An optimized design minimizes background and maximizes spot intensity by using many high-affinity probes, resulting in high SNR and improved detection performance.

Despite its importance, smRNA-FISH probe design is often underappreciated. Probe design has two facets: (i) the *physical* signal amplification strategy design, which constructs labeling strategies to enhance or amplify the fluorescent signals that arise from single-stranded DNA probes binding RNA transcripts (18), and (ii) *computational* design, which selects optimal nucleotide sequences for hybridization (19). High-fidelity design must consider numerous variables that influence performance, including oligonucleotide thermodynamics, hybridization conditions, genome complexity, probe secondary structure, gene expression variability, and genomic coverage (19).

Commonly used state-of-the-art software—Stellaris (20), MERFISH (15), Oligostan-HT (21), and PaintSHOP (22)—share a similar workflow (Figure S1) but apply different criteria. Common limitations include narrow default windows for melting temperature (T_m_) and GC content, incomplete quantification of multi-probe off-target binding, and—except for MERFISH—little integration of expression data that may modulate off-target signal. Consequently, designs may return too few probes for atypical genes or probes prone to false positives.

These constraints are particularly problematic for short genes ≤ 2kb (insufficient probe numbers), low-abundance transcripts (signal overwhelmed by background), genes with tissue-specific expression (variable performance), sequences with shared motifs (cross-hybridization), and low-complexity or lncRNA targets (excess off-targets).

To overcome the issues observed with other probe-design software, we developed TrueProbes, a design platform that models genome-wide binding affinities to create application-specific, high-specificity probe sets. By evaluating off-target interactions genome-wide, TrueProbes seeks to more accurately minimize off-target mediated background fluorescence (Figure 1A). In this approach, we conceptualize the number of probes and their on-/off-target binding probabilities to jointly determine on-target spot intensity (dark green) and off-target background (light green); reducing their overlap improves true RNA detection (Figure 1B).

TrueProbes operates via the command line in MATLAB or as a standalone application using MATLAB Runtime on macOS, Windows, and Linux and can be tailored to diverse applications: mature-RNA detection, intronic nascent RNA detection, isoform-specific or agnostic designs, cell- or tissue-specific panels, and multi-fluorophore labeling (see TrueProbes manual for instructions on tailoring probe design to specific applications). The software simulates probe performance across probe/salt concentrations, hybridization temperatures, kinetic models, and user-provided expression datasets, and it can refine existing probe sets by removing sequences predicted to impair performance. Collectively, these computational advances translate into probe sets that exceed the experimental performance of current smRNA-FISH tools.

### Computational simulation of probe performance demonstrates and highlights the strengths and weaknesses of each software’s algorithms

Target‐gene expression fundamentally shapes smRNA-FISH performance. When a transcript is highly expressed, individual RNA molecules often appear so close together that resolving single spots becomes difficult; at the other extreme, very low expression requires imaging many cells to capture only a few spots. Therefore, any reduction in on-target gene expression lowers the average number of detectable spots per cell, even though the signal-to-noise ratio of each spot remains comparable due to probe-to-target binding kinetics remaining unchanged (Figure 1C).

Probe specificity can be assessed in knockout (KO) cells by monitoring changes in background intensity attributable to off-target binding (Figure 1D). Interpreting KO data, however, can be complicated by compensatory shifts in the expression of the probes’ off-target genes.

TrueProbes aims to approximate the ideal scenario in which probes bind their intended targets strongly and possess only weakly bound off-targets, thereby maximizing signal-to-noise (Figure 1E).

Most existing design programs generate probes sequentially from the 5′ to 3′ end of the transcript instead of ranking candidates globally by experimental utility (Figure 2A). A common workflow tiles the RNA with all oligos that meet length constraints, then filters them according to each tool’s heuristics (Figure 2B). Stellaris, Oligostan-HT, PaintSHOP, and MERFISH all follow a classic three-step path—filter, evaluate off-targets, and select a non-overlapping set—but they diverge in how they define and score specificity (Figure S1).

**Figure 2:**
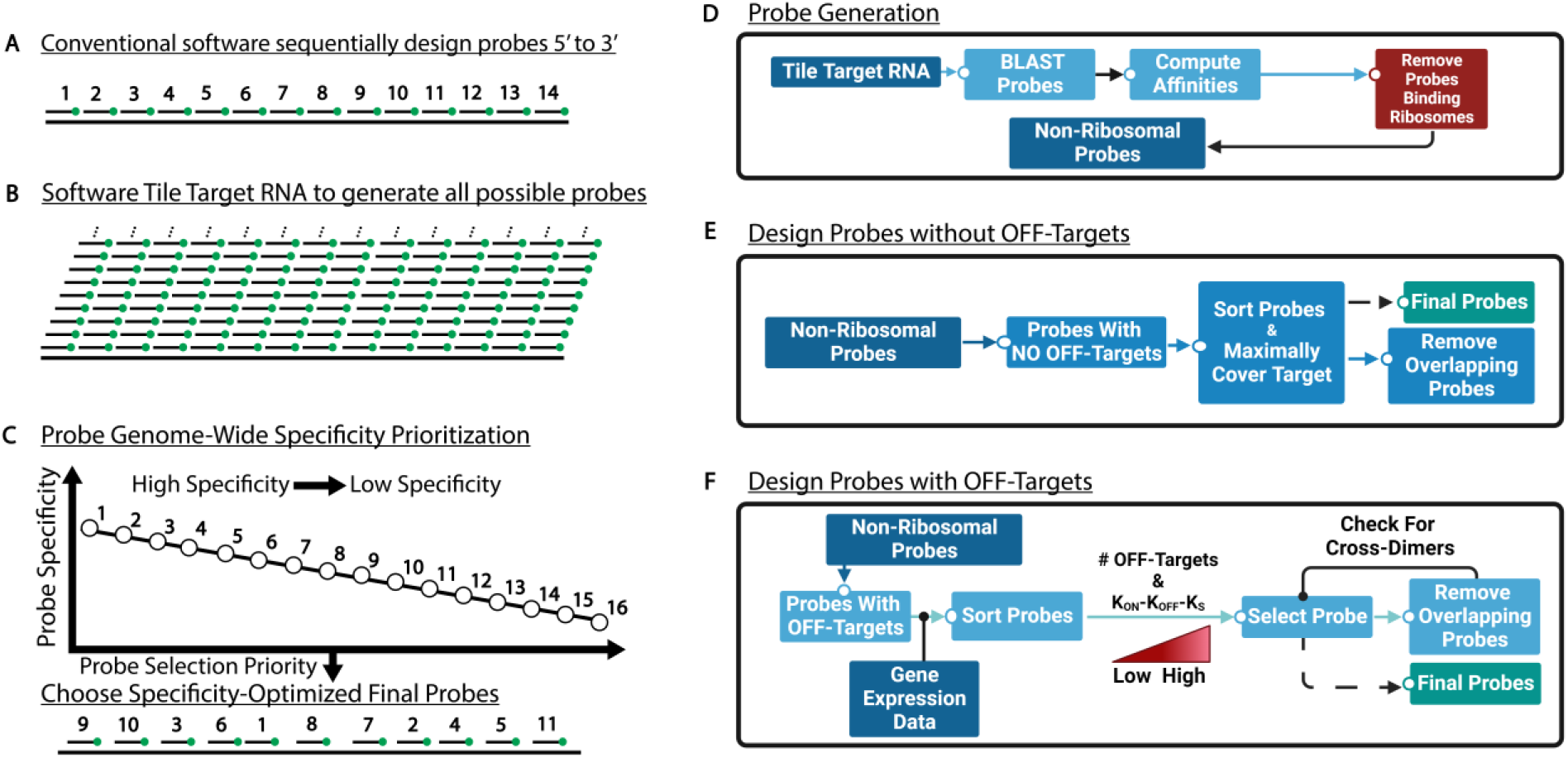
TrueProbes systematically evaluates on- and off-target binding dynamics to generate probe sets optimized for individual experimental contexts. (A) Traditional smRNA-FISH design tools often select probes sequentially from the 5′ to 3′ end of the target transcript, introducing bias based on probe position rather than optimizing for binding strength or specificity. (B) Most software platforms tile the RNA with candidate probes of defined length, then apply heuristic filters to select probes, without global ranking. (C) In contrast, our approach ranks all candidate probes based on experimentally relevant specificity scores, prioritizing probes with strong on-target binding and minimal off-target interactions. (D) The pipeline begins by tiling the full target RNA sequence and generating all possible probe sequences. Each candidate is screened for off-target binding events of ≥15 nt using BLAST, assessed for RNA secondary structure, and evaluated for thermodynamic stability, including hairpins, cross-dimers, and rRNA interactions. Probes that bind rRNA are excluded. (E) Among probes with no detectable off targets, the software selects those that provide maximum coverage of the target RNA, ranking candidates by corrected on-target binding affinity after accounting for RNA secondary structure. (F) For probes with off targets, with or without incorporating cell-type–specific expression data, the remaining candidates are ranked by their specificity (low off-target binding, strong on-target binding, and minimal self-structure). The final probe set is assembled by iteratively selecting top-ranked probes that do not cross-dimerize with one another.

Stellaris first removes 17–22-mer candidates outside a narrow GC-content window, then sweeps 5′ to 3′ while applying five masking levels to discard repetitive or non-species–specific sequences; the first oligo that satisfies length, spacing, and T_m_ criteria is accepted, yielding a largely “first-pass” design (Figure S1A). MERFISH also filters on GC/T_m_, but hashes each oligo into 15/17-mers and compares them against the transcriptome and rRNA to compute an off-target index; probes are retained only if RNA scores are ≤ 0.7 and rRNA scores are 0, followed by a greedy 5′ to 3′ spacing step (Figure S1B). Oligostan-HT applies GC and low-complexity screens, then calculates Gibbs free energy (ΔG°) for every remaining oligo, ranking each by proximity to a user-defined optimum and selecting the highest scores until the quota is met (Figure S1C). PaintSHOP combines conventional thermodynamic filters with a Bowtie2 alignment stage to eliminate oligos with many genomic matches; a machine-learning (ML) classifier then assigns each candidate the probability of forming deleterious off-target duplexes, and only the top-ranked, non-overlapping probes are kept (Figure S1D). Collectively, these methods span a spectrum from heuristic, position-ordered choices (Stellaris) through energy-based ranking (Oligostan-HT), alignment plus ML triage (PaintSHOP), to hash-based transcriptome screening (MERFISH).

TrueProbes instead ranks all candidates from highest to lowest predicted specificity—defined by minimal expressed off-target binding, strong on-target affinity, weak off-target affinity, low self-hybridization, and minimal cross-dimerization—before assembling the probe set (Figure 2C). TrueProbes first uses BLAST to enumerate off-targets, removes probes that bind to rRNA, and then calculates on-target and off-target binding energies for every oligo (Figure 2D). Probes with no off-targets are selected first, maximizing coverage while excluding sequences with significant cross-dimerization to those already chosen (Figure 2E). Remaining candidates are ranked by (i) the number of off-targets—optionally weighted by gene-expression data—and (ii) the difference of on-target to off-target plus self-hybridization energies; top-ranked, non-overlapping probes are added iteratively (Figure 2F).

For every probe set, TrueProbes can simulate expected smRNA-FISH outcomes—including optimal probe, RNA, and salt concentrations—and optionally account for probe secondary structure, hybridization temperature, multiple targets, fluorophore choice, DNA, nascent RNA, and photon-count statistics (Figures S2A–S2B). The model can be used to generate predictions for temperature and cell-line sensitivity, multi-target discrimination, multiple fluorophore colocalization; when provided transcript expression levels and probe/background intensity, it can start to generate predictions for spot intensity, background, signal-to-noise ratio, and false-positive/-negative rates (Figure S2C). Additionally, for smRNA-FISH probe sets, TrueProbes can identify with or without gene expression data which probes contribute the most to i) the total number of off targets or ii) the accumulation of off targets with two or more binding sites bound by probes.

To benchmark performance, we searched genome-wide for transcripts that challenge conventional probe design, including due to certain attributes—short length, low or variable expression, complex isoform architecture, shared sequence motifs, low complexity, or overlapping antisense lncRNAs—and then chose genes meeting multiple criteria (Figures S3A– S3B).

As a representative example, we selected ADP-ribosylation factor 4 (ARF4), a small GTPase implicated in epidermal growth-factor signaling, COPI vesicle formation, recycling-endosome integrity, ciliogenesis, and several disease processes, including cancer progression and cholera-toxin activation (23). In addition to its biological and medical importance, ARF4 was selected for final experimental validation because it had the highest gene expression (in TPM units) and the fewest isoforms among all candidate genes for the Jurkat cell line.

For ARF4, TrueProbes, Stellaris, Oligostan-HT, PaintSHOP, and MERFISH designed 50, 51, 29, 17, and 18 probes, respectively (Figure 3A). We computationally assessed cumulative off-target probe binding under various assumptions using RNA-seq data expressed in Trimmed Mean of M-values (TMM)-normalized transcripts per million (nTPM) (Figure 3B–G) (24).

**Figure 3:**
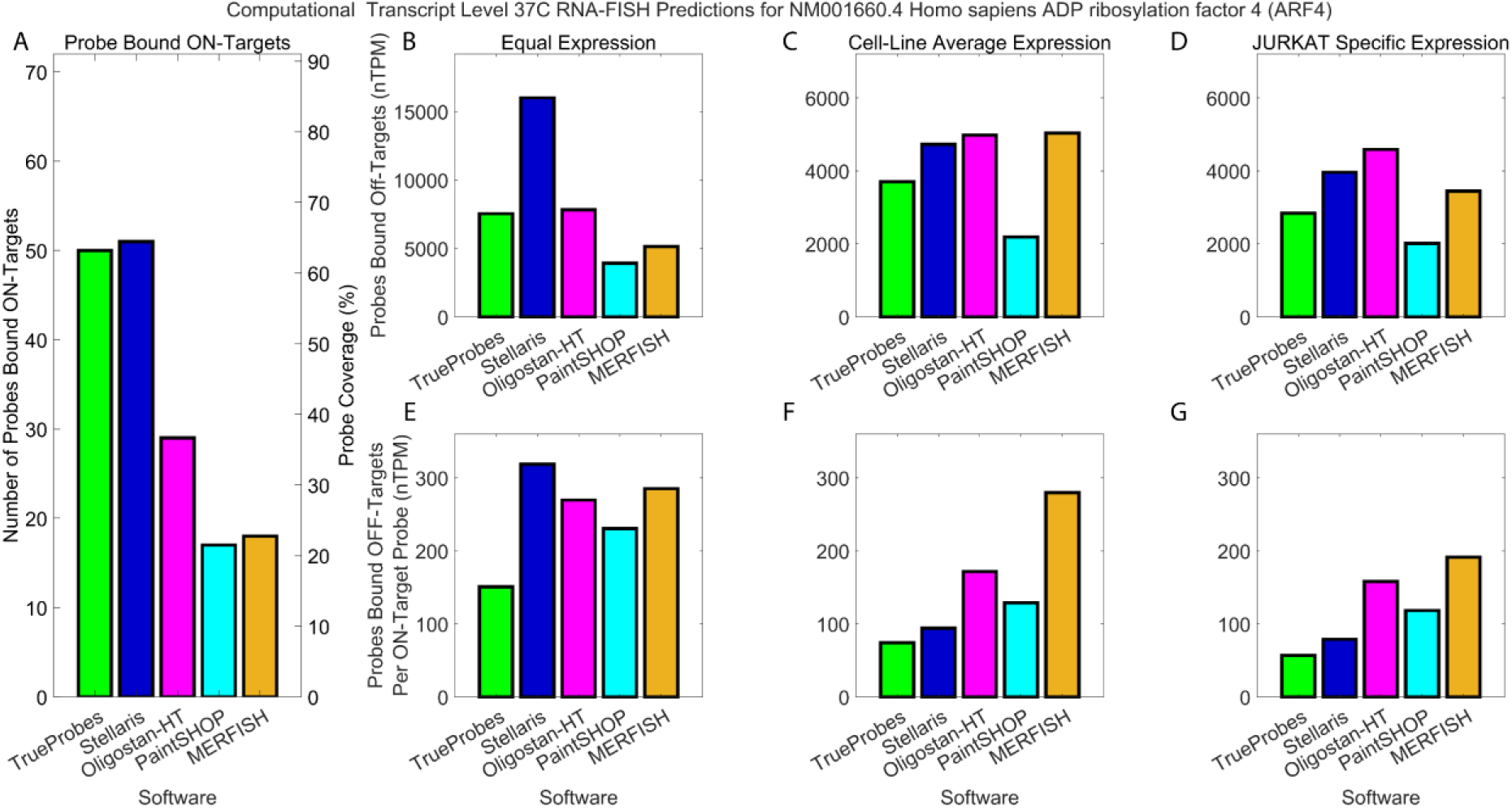
Systematic probe design improves signal and reduces off-target background. (A) Absolute number of probes designed for *ARF4* (left) and percentage of gene coverage (right) for each software: TrueProbes (green), Stellaris (blue), Oligostan-HT (pink), PaintSHOP (cyan), and MERFISH (orange). (B–D) Cumulative number of probes predicted to bind off-target RNAs at equilibrium under three transcript expression assumptions: equal expression across genes (B), average cell line expression (C), and Jurkat-specific transcript expression (D). (E–G) Ratio of off-target to on-target probe binding events under the same conditions, quantifying specificity of each design. Panels highlight the advantage of expression-aware, thermodynamically optimized probe design in minimizing background and maximizing true signal.

Under the assumption of equal transcript expression across all genes—the current standard for smRNA-FISH probe design—Stellaris exhibited the highest number of probes bound to off-targets (Figure 3B). In comparison, TrueProbes and Oligostan-HT showed approximately half as much total off-target binding as Stellaris. PaintSHOP and MERFISH exhibited the lowest off-target accumulation, approximately 50–75% lower than that of TrueProbes and Oligostan-HT (Figure 3B).

When accounting for gene expression differences using mean average expression levels across cell lines, the total number of off-target probe binding events decreased for all software tools due to many off-targets not being expressed in several cell lines (Figure 3C). Stellaris, Oligostan-HT, and MERFISH had the highest overall off-target binding in this scenario. TrueProbes showed approximately 20% fewer off-target bindings than these three, while PaintSHOP reduced off-target accumulation by ∼40% compared to Stellaris, Oligostan-HT, and MERFISH (Figure 3C).

Using Jurkat cell-specific expression data, Oligostan-HT had the highest total off-target binding, followed by Stellaris and MERFISH (Figure 3D). TrueProbes and PaintSHOP showed approximately 40% and 60% fewer off-target bindings, respectively, compared to Oligostan-HT (Figure 3D).

To assess probe specificity, we normalized off-target binding by the number of probes designed by each software. Under equal transcript expression, Stellaris, Oligostan-HT, PaintSHOP, and MERFISH had comparable off-target accumulation rates per probe (Figure 3E). In contrast, TrueProbes showed a substantially lower rate of off-target binding per probe designed (Figure 3E).

When using cell-line-averaged mean expression, TrueProbes and Stellaris both accumulated about 75 and 90 off-targets per on-target binding event, respectively (Figure 3F). Oligostan-HT and PaintSHOP exhibited roughly double the number of off-targets per on-target (150, 120) compared to TrueProbes and Stellaris. MERFISH’s performance remained consistent with its performance under the equal-expression scenario, with approximately three to four times more off-targets per probe compared to TrueProbes and Stellaris (Figure 3F).

Finally, using a kinetic equilibrium model to estimate the mean number of probes bound on target RNA in Jurkat cells, we calculated probes bound off targets per on-target designed probe. MERFISH had the highest off-target-to-on-target ratio, followed by Oligostan-HT, PaintSHOP, Stellaris, and TrueProbes (Figure 3G). Similar trends were observed when using gene-level expression data instead of transcript isoform-resolved data, with only slight variations in off-target binding (Figure S4A–S4G).

### RNA-FISH Experimental Results Demonstrate that Off-Target and Binding Affinity Inclusive Probe Design Improve RNA-FISH Signal Discrimination

To assess the efficacy of RNA-FISH probes in cell culture, the probe for hARF4 coupled to Cy5 from each design tool was imaged in Jurkat cell cultures. Representative images show the presence of smRNA-FISH spots (green) in DAPI-stained (blue) Jurkat cells when hybridized without probes (Figure 4A) and with smRNA-FISH probes from TrueProbes (Figure 4B), Stellaris (Figure 4C), Oligostan-HT (Figure 4D), PaintSHOP (Figure 4E), and MERFISH (Figure 4F). Images from all biological replicates were then processed through the same pipeline: spot localization, intensity extraction, and cell-specific thresholding. For each cell, we recorded the maximum pixel value of each spot (signal), the mean local background, the signal-minus-background value, the signal-to-noise ratio (SNR), and the total number of detected spots. Uneven illumination was corrected with the blank-control images, and replicate-specific thresholds were applied.

**Figure 4:**
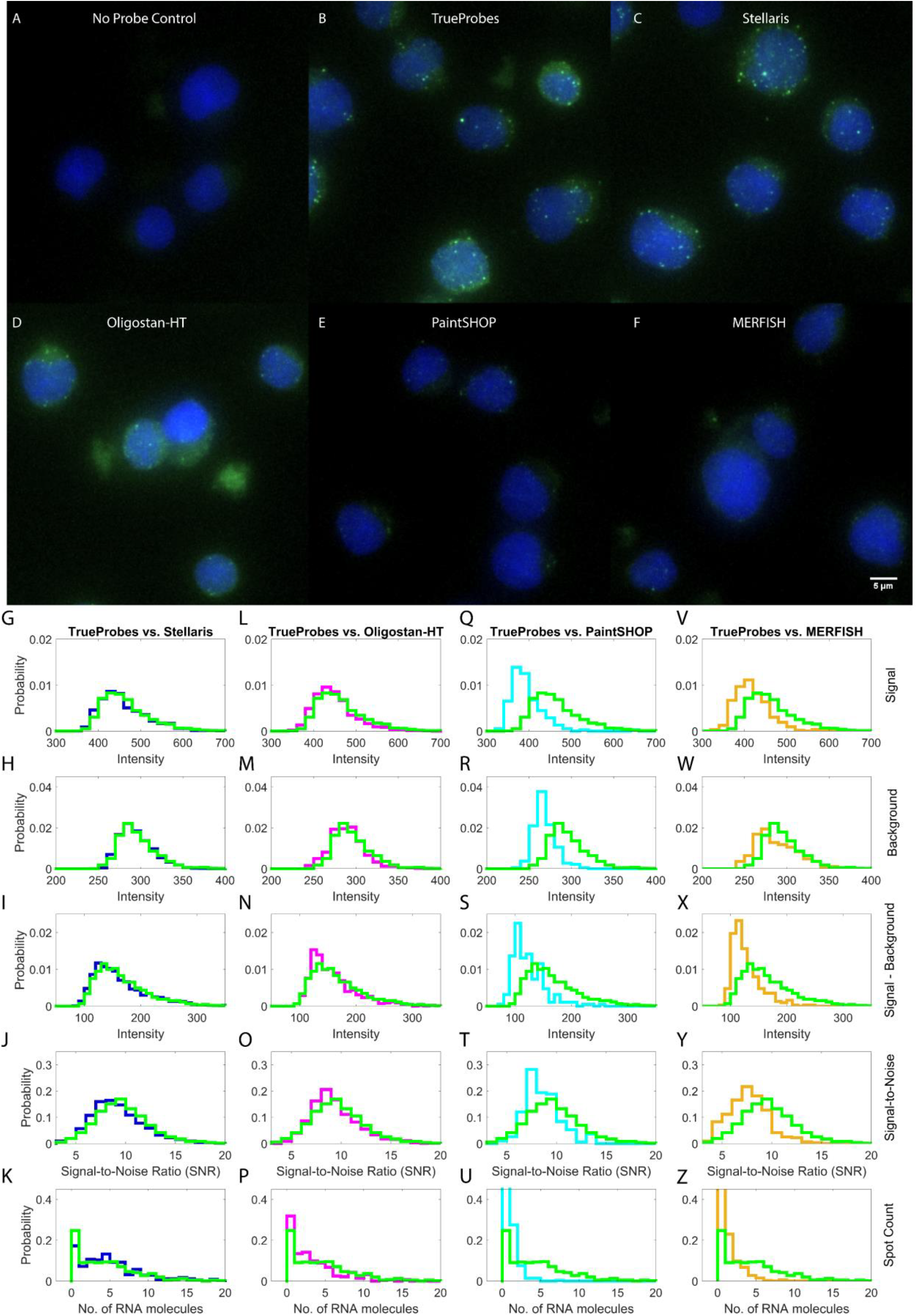
Experimental validation of smRNA-FISH probe designs for ARF4. Representative Jurkat cell field of view with ARF4 RNA spots (green) and DAPI staining (blue) for no probe control (A), TrueProbes (B), Stellaris (C), Oligostan-HT (D), PaintSHOP (E), and MERFISH (F). Distributions are shown for spot signal intensity (G, L, Q, V), background intensity (H, M, R, W), signal-minus-background (I, N, S, X), signal-to-noise ratio (J, O, T, Y), and single-cell spot counts (K, P, U, Z). Each row corresponds to a different design tool: Stellaris (G–K, blue), Oligostan-HT (L–P, magenta), PaintSHOP (Q–U, cyan), and MERFISH (V–Z, beige), with all compared to TrueProbes (green) across three experimental replicates. TrueProbes: 409 cells, 1841 RNA spots; Stellaris: 348 cells, 1753 spots; Oligostan-HT: 283 cells, 1,014 spots; PaintSHOP: 291 cells, 191 spots; MERFISH: 327 cells, 314 spots.

Probe sets designed with TrueProbes, Stellaris, and Oligostan-HT produced nearly identical distributions for spot signal intensity (Figure 4G, L), background intensity (Figure 4H, M), signal-minus-background (Figure 4I, N), SNR (Figure 4J, O), and per-cell transcript counts (Figure 4K, P). In contrast, probes from PaintSHOP and MERFISH yielded lower overall performance, whereas TrueProbes retained higher median values for signal intensity relative to PaintSHOP and MERFISH (Figure 4Q, V), background (Figure 4R, W), signal–minus–background (Figure 4S, X), SNR (Figure 4T, Y), and spot counts (Figure 4 U, Z). Replicate-to-replicate variability was observed, but the relative ranking of probe sets remained consistent across experiments (Figure S5). Interestingly, the probes designed using PaintSHOP and MERFISH resulted in almost double the number of non-expressing cells in comparison to other probe design platforms (Figure 4K, P, U, Z).

## Discussion

Accurate quantification of single-cell gene expression critically depends on the specificity and sensitivity of probe design in smRNA-FISH experiments. Our results demonstrate that current design platforms vary widely in their ability to minimize off-target binding and maximize true signal, particularly when targeting challenging transcripts such as *ARF4*, a gene with clinical and mechanistic relevance.

Our computational analysis highlights several key trends. First, expression-aware design significantly improves probe specificity. Designs assuming equal transcript abundance—a common simplification—consistently overestimated off-target binding across all tools. By incorporating average or cell-line–specific gene expression, total off-target binding dropped substantially, suggesting that realistic expression models are essential for high-fidelity design. This improvement is particularly important for transcripts expressed in a cell–type–specific or dynamic manner, where misestimated background can compromise detection accuracy.

Among the tested tools, TrueProbes consistently demonstrated the lowest off-target accumulation per probe and the most favorable signal-to-noise predictions across multiple expression scenarios. This advantage stems from its genome-wide modeling of RNA-probe binding kinetics, integration of cell-type-specific transcriptomic data, and a global specificity-based ranking system that accounts for thermodynamic stability, self-dimerization, and cross-hybridization. Notably, TrueProbes computes site-specific binding probabilities and kinetic binding affinities across the transcriptome, enabling precise estimation of both on-target and off-target interactions.

By contrast, MERFISH and PaintSHOP produced probe sets that exhibited relatively poor signal and elevated off-target accumulation. This disconnect highlights the importance of not only assessing potential binding sites but also integrating binding strength, transcript abundance, and cross-reactivity into a unified performance metric.

Despite generating a similar number of probes, Stellaris consistently demonstrated higher off-target interactions and greater background signal compared to TrueProbes. Oligostan-HT, on the other hand, used an affinity-based greedy method that resulted in slightly smaller probe sets than TrueProbes, but still had higher off-target binding and greater background signal. These findings for Stellaris and Oligostan-HT together show that while masking out repetitive sequences and only focusing on probe on-target binding can make it easier to design larger probe sets, it is not enough to fully optimize probe performance to avoid specific off-targets that are non-repetitive in sequence. This further highlights that probe off-target information needs to be included in the probe design process to lower the risk of performance variability when target genes share some amount of sequence homology with others in the target genome.

Experimental validation supported the observed performance trends. Probes grouped into two distinct performance tiers. Group 1—TrueProbes, Stellaris, and Oligostan-HT—showed comparable results across key metrics, including signal intensity, signal-minus-background, signal-to-noise ratio, and spot counts. In contrast, group 2—PaintSHOP and MERFISH—underperformed across all major metrics, exhibiting lower signal intensity, lower spot counts, and reduced SNR. They also showed a higher fraction of cells lacking detectable RNA expression, likely due to limited probe coverage or increased off-target interference from cell-specific expression of off-target sequences.

Probe length is another factor crucial for providing context on how RNA-FISH software probe sets performed. The software’s used a variety of probe length ranges with TrueProbes and Stellaris using short probes (20nt), Oligostan-HT using a mixture of short and long probes (30.1 ± 2.4 nt, range 25 – 34 nt, for *ARF4* from 23 – 38 nt probe tiling), PaintSHOP using longer probes (33.8 ± 3.2 nt, range 30 – 34 nt, for *ARF4* from 30 – 37 nt probe tiling), and MERFISH using longer probes (30nt). Choosing the optimal probe length can be a delicate balance as increasing probe length, i) decreases the likelihood of observing full sequence matches; ii) increases probe binding affinity and the likelihood it will bind targets; iii) increases secondary structure binding affinity and reduces the likelihood it will bind targets; and iv) for short genes decreases the number of probes you can design and lowers spot detection.

Previous work (25, 26) has given some justification for using longer probes, finding that when assessing reference genomes that short DNA sequences (20 – 25 nt) can have higher frequencies of full repeated sequence matches that can result in lower probe specificity and provided a basis for using k-mer/hash counting methodology in probe design, included in PaintSHOP and MERFISH. This, however, does not show how easy it is to find a subset of short probes with few off-targets. While it is true that increasing probe length does lead to, on average, stronger binding affinity and increased sequence uniqueness lowering chances of full sequence matches, our computational predictions and experimental results show that i) even shorter probes we find can have enough binding affinity to bind on-target with near 100% probability; ii) longer probes subsequence off-target sequence matches that still have enough complementary to bind off targets with near 100% binding probability. These two points together show that with careful off-targets and binding equilibrium, inclusive probe design can create short probes that outperform longer probes.

Furthermore, our predictions also provide evidence about the usefulness of avoiding probe secondary structure during RNA-FISH probe design. Our computational modeling also shows that when combining probe secondary structure and probe-target binding into a binding comprehensive equilibrium metric, most probes’ secondary structure is predicted to have minimal impact on probe binding compared to the probe on-/off-target binding affinities, especially as probes are often in far excess of targets. This effect, though, might be underestimated given how software already tries to avoid secondary structure in the probes they design.

Our results also have implications for improving how we design oligonucleotide therapies, where oligos are designed to either degrade RNA or sterically inhibit mRNA translation in disease treatment (27). Probe selectivity and specificity are critical to i) improving therapy sensitivity by increasing target binding affinity and knockdown (KD) efficiency; and ii) mitigating off-target effects from KD of probe off-targets by lowering the number of probes off-targets and probe off-target binding affinities.

Another potential application of TrueProbes is in designing antisense oligonucleotides with reduced off-target binding, enabling more specific RNA capture for downstream analysis (28).

Looking ahead, TrueProbes’ capacity to integrate expression data into the off-target evaluation process provides a strong foundation for designing cell-line and tissue-agnostic probe sets, in which probe performance remains stable across varying expression contexts. This is especially relevant for applications such as multiplex smRNA-FISH and spatial transcriptomics (18), where probes for multiple genes are used simultaneously. In these cases, the off-target burden may compound across genes, and probes that perform well individually may fail in combination. Our findings emphasize the need for the preselection of minimally overlapping genes to reduce cumulative background in multiplex assays.

Collectively, TrueProbes demonstrates how a first-principles approach—grounded in physical modeling, expression-aware off-target analysis, and comprehensive ranking—can overcome many of the limitations seen in current smRNA-FISH design tools. By improving signal-to-noise and expanding probe accessibility for difficult targets, TrueProbes enables broader and more reliable use of single-cell RNA imaging technologies.

### Assumptions and Limitations

The TrueProbes smRNA-FISH probe design software and underlying kinetic model rely on a set of assumptions, driven primarily by the limited availability of quantitative data on factors that can influence smRNA-FISH performance. In cases where these assumptions are not valid, the predictive accuracy of the software may decline. For example, the model assumes that off-target transcripts are uniformly distributed throughout the cell, but RNA subcellular localization can introduce spatial variability in the background signal. Off-target accumulation in specific compartments may lead to localized increases in false-positive spots if multiple off-targets colocalize. The model also focuses exclusively on fully spliced transcripts and does not account for binding to intronic regions, meaning that nascent transcriptional dynamics, including on-target or off-target binding at transcription sites, are not captured. This limits the accuracy of predictions for nuclear-localized RNA signals. RNA secondary and tertiary structures are not included, which may lead to inaccuracies if binding sites are structurally occluded. Similarly, RNA–protein interactions, which can modulate accessibility of the transcript, are not modeled. Probe diffusion kinetics are also not simulated—we assume that sufficient time has passed during the experiment for probes to distribute throughout the cell and reach equilibrium binding states (29). Additionally, the model does not include toehold-mediated strand displacement (TMSD), a process by which one probe may displace another from the same target RNA, potentially affecting equilibrium probe occupancy and the rate of RNA–oligo hybrid formation (30). Partial binding of overlapping probes is likewise not considered, as we do not model intermediate states of sequential probe binding (31). Formamide concentration, which modulates the thermodynamic stability of probe–RNA interactions and disrupts DNA/RNA duplexes (32), is also not explicitly included. Additionally, all probe-target binding affinity calculations were made after filtering out sequence mismatches in probe-target paired hybridization sequences, which underestimates off-target binding in cases where mismatches can contribute to binding affinity that rival on-target binding affinity (33). This may impact predictions of probe binding strength, especially in cases where mismatches form energetically stable interactions that mimic full sequence complementarity. We currently include a mesoscopic model of probe-target mismatch binding (33), which is not evaluated by default due to having much longer runtimes than using nearest-neighbor Gibbs free energy models. We have since added a nearest-neighbor model approximation of probe mismatch hybridization Gibbs free energy (34) to the models that TrueProbes can use to evaluate probe binding affinities. Our off-target predictions are based on a reference genome and transcriptome, and any discrepancies or incompleteness in these references could affect design outcomes. Moreover, the variability in gene expression between cells, conditions, or cell types presents inherent limitations. In certain single-cell expression contexts, designing a probe set that minimizes off-target binding below a threshold acceptable across all cells may be theoretically impossible. The same challenge applies across experimental conditions, where a probe set optimized for one context may not retain its performance in others. Finally, genes with high sequence similarity or shared motifs may produce smRNA-FISH signals that are experimentally indistinguishable, particularly under noisy conditions, limiting the ability to assign detected spots to a specific transcript confidently.

### Experimental Methods

#### smRNA-FISH Probe Creation

Probe DNA oligos with 3’ amine modifications were ordered from LGC Biosearch Technologies. smRNA-FISH probe sets were made by conjugating DNA oligos to Amersham™ CyDye™ Cy5 Mono-Reactive Dye (PA25001), followed by HPLC purification. A SpeedVac Vacuum concentrator was then used for 8 to 12 hours to pool and resuspend combined HPLC-purified probe sets in 50 µL of nuclease-free water. Conjugated oligo-CY5 probe set concentrations were quantified using an Implen VU-VIS Spectrometer NanoPhotometer P300.

#### Cell-Culture

Jurkat cells were grown from individual vials by expanding one vial of cells. Jurkat media was made by adding 10% heat-inactivated Fetal Bovine Serum (FBS) (Corning 161140-071), 1% 100x Penicillin-Streptomycin (Gibco 15140-122), and 1% 100x GlutaMax (Corning 35050-061) to 500 mL of RPMI 1640 (Corning 15-040-CV). The initial vial was grown in Jurkat media for several days to get about 15 million.

#### Cell-Fixation

The Jurkat cells were fixed for initial smRNA-FISH experiments and probe concentration optimization. Jurkat cells were washed by centrifuging for 5 minutes at 300xg and resuspending the pellet in 1 mL 1x phosphate-buffered saline (PBS). They were then transferred to 1.5 mL LoBind® tubes. Cells were centrifuged for 5 minutes at 300xg, and the supernatant was removed. Cells were then resuspended in 1 mL fixation buffer (1x PBS, 3.7% formaldehyde) and, after mixing, incubated at room temperature for 10 minutes. Cells were centrifuged for 5 minutes at 300xg and washed twice with 1 mL PBS to remove formaldehyde. Cells were finally resuspended in 1 mL 70% ethanol overnight and stored at 4°C until needed.

#### DAPI and smRNA-FISH

BSA-coated tips were used to transfer cells into LoBind® tubes. Cells were then centrifuged at 300xg for 5 minutes, and ethanol was aspirated. The fixed cells were equally divided into six experimental groups resulting in more than 2million cells per group, one group as no probe control and five groups one for each design tool. Cells were resuspended in 100 µl wash buffer (1x PBS with 10% formamide) and centrifuged for 5 minutes at 300xg. Wash buffer was removed by aspiration, and then cells were resuspended in 50 µl of hybridization buffer, which consists of 10% dextran (Sigma Aldrich, D8906), 10 mM vandyl-ribonucleoside complex (NEB, S1402S), 0.02% RNAse-free Bovine Serum Albumin (BSA) (Ambion, AM2616), 1 mg/mL *E. coli* tRNA (Sigma, R4251), 2x saline-sodium citrate (SSC) (Invitrogen, 15557-044), 10% formamide (Invitrogen, 15515-026), and smRNA-FISH probes for hARF4 at molar-adjusted dilution from stock concentration. Probe set concentrations were diluted to have the same final equal molar concentration per probe in each probe set as the TrueProbes 1:2000 probe set stock dilution. Cells were added to hybridization buffer and then incubated overnight at 37°C. The next day, cells were washed once with 1 mL wash buffer and resuspended in 1 mL wash buffer with 0.5 µg/mL DAPI (Sigma, D9542-10MG). Cells were then incubated for 30 minutes at 37°C, followed by centrifugation, aspiration of wash buffer with DAPI, and transferred to wash buffer without DAPI. Cells were stored at 4°C until imaging.

#### Microscopy

Micro-Manager software version 1.4 was used to control a Nikon Ti-2 Eclipse epifluorescence microscope under perfect focus (Nikon), using a 100x VC DIC lens (Nikon), X-cite Xylis LED fluorescent light source (Excelitas), and an Andor Sona-11 sCMOS Camera with a pixel size of 11 µm (Oxford Instruments). Cells were mounted in a 15 μ-Slide Angiogenesis slide (ibidi GmbH, 81506) in wells coated with 0.01% poly-D-lysine (Cultrex 3439-100-01). Fluorescent signals were imaged using Semrock Brightline filters for DAPI (DAPI-5060C-NTE-ZERO) and CY5 (NIK-0013 RevA-NTE-ZERO). Jurkat cells were imaged with epifluorescence and transillumination at 100x magnification with 500ms CY5 exposure time, 100ms TRANS exposure time, and 50ms DAPI exposure time, at 300nm per z-step.

### Computational Methods

#### Reference Genome and Gene Expression Data Acquisition

Genome and transcriptome reference assemblies were obtained from the Genome Reference Consortium (GRC) in addition to NCBI and EMBL-EBI gene annotations for *Danio rerio* (danRer11; GCF_049306965.1, GCA_000002035.4), *Saccharomyces cerevisiae* (sacCer3; GCF_000146045.2, GCA_000146045.2), *Mus musculus* (mm39; GCF_000001635.27, GCA_000001635.9), and *Homo sapiens* (hg38; GCF_000001405.39, GCA_000001405.29) (35, 36). Additional reference sequence metadata—such as transcript identifiers, chromosomal positions, and exon annotations—was collected from NCBI RefSeq (35), the EMBL Database (36), and the GENCODE Project (37). Preprocessed expression profiles from bulk RNA-sequencing studies: the Cancer Cell Line Encyclopedia (CCLE) (38), the Genotype-Tissue Expression (GTEx) Project (39), the Cancer Genome Atlas (TCGA) (40), the Human Protein Atlas (41); and large-scale single-cell RNA-sequencing studies (42, 43) are integrated into TrueProbes’ database data available to be used for designing RNA-FISH probes. For Human Protein Atlas cell lines, lncRNA expression was imputed using the average lncRNA-to-mRNA expression ratio in corresponding tissue sources (39, 40).

#### smRNA-FISH Probe Design Target Gene Selection

The list of candidate genes for experimental smRNA-FISH validation was selected using bulk RNA-seq data and gene sequence information from the Human Protein Atlas (41). All chosen mRNAs were between 500 and 3000nt long, had fewer than five isoforms, had enough space for at least twenty isoform-specific and agnostic probes, and had antisense lncRNA between 500nt and 3000nt. Selected genes were also required to have at least an average expression of 100 transcripts per million (TPM). Using cutoffs for each selection criterion, we obtained twenty-nine final gene isoforms. The genes include sense/antisense RNA for (MRPL20, IGFBP8, NOP53, TM4SF1, ARF4, RAB5C, PRDX6, YWHAH, VIM) as well as positive control genes GAPDH and ACTB.

#### Stellaris Probe Designer Parameters

Probes were designed using the Stellaris Probe Designer version 4.2 (https://www.biosearchtech.com/stellaris-designer) with a probe length of 20 nucleotides and a minimum spacing of 3 nucleotides between adjacent probes, using masking levels 1 through 5. Probe sets from masking level 3 were selected, as this level consistently yielded the highest number of probes when aiming to design at least 24 or 48 probes. Masking levels 3–5 apply species-specific filters to avoid cross-hybridization with RNAs that Stellaris identifies as commonly or highly expressed.

#### Oligostan-HT Probe Designer Parameter Settings

Probes were designed using the Oligostan-HT tool, acquired via <u>Docker</u> (https://hub.docker.com/r/oligostan/oligostan_ht_rna). Probe lengths were set to the default range of 23–38 nucleotides, and minimum spacing between adjacent probes was adjusted to 3 nt. Repetitive sequences were filtered using RepeatMasker with a maximum masked percentage threshold of 10%. Binding affinity was scored based on the deviation of each probe’s RNA-RNA hybridization Gibbs free energy (ΔG°) from the target ΔG° value of -46 kcal/mol at 48°C, using a normalized energy score. Only probes with energy scores between 0.7 and 0.99 were retained. For each candidate gene isoform, probes were first designed using Oligostan-HT Design #1 (GC-content 35–65%); if no probes were identified, Design #2 (GC-content 25–70%) was used. The default minimum and maximum number of probes per target were reduced from 50 to 1 and 100 to 96, respectively. Designs #3 and #4—which permit probes with overlapping positions— were excluded to maintain consistency with non-overlapping probe design principles. All other nucleotide composition and stacking filters were retained at default settings kept at false.

#### PaintSHOP Probe Designer Parameter Settings

The PaintSHOP pipeline source code (https://github.com/beliveau-lab/PaintSHOP_pipeline) and isoform-resolved probe sets for the hg38 genome (https://github.com/beliveau-lab/PaintSHOP_resources) were downloaded from GitHub. Probes were generated as described previously (22), using repeat-masked regions, default probe lengths of 30–37 nt, and a melting temperature range of 42–47°C. The design used default 0-nt spacing between probes, GC-content between 20–80%, and a thermodynamic model based on DNA/DNA nearest-neighbor rules (44). Additional constraints included a maximum self-homology of 6 nt, cross-homology of 8 nt, a Na^+^ concentration of 390 mM, and 50% formamide. Hybridization scores were computed at 42°C using PaintSHOP’s XGBoost-based machine learning model. Probes containing five-nucleotide homopolymer runs (‘AAAAA’, ‘TTTTT’, ‘CCCCC’, or ‘GGGGG’) were excluded.

#### MERFISH Probe Designer Parameter Settings

MERFISH probe design was performed using code from both the Zhuang Lab (https://github.com/ZhuangLab/MERFISH_analysis) and the Allen Institute (https://github.com/AllenInstitute/MERFISH_analysis) obtained from GitHub. The Allen Institute version was modified to incorporate 2021 updates from the Zhuang Lab, enabling barcode generation at the transcript isoform level rather than the gene level. Probes were fixed at 30 nt in length with 3-nt spacing. Probe melting temperature requirements were maintained in the default range of 70–100°C, using the DNA/DNA nearest-neighbor free energy model (45) with 300 mM Na^+^. GC-content constraints were kept at 0–100%. Specificity scoring used hash tables to evaluate exact 17-nt matches (for isoform specificity) and 15-nt matches (for rRNA penalties). Default specificity thresholds were applied (0–1.0 for isoform-specific and 0.7–1.0 for general specificity). Probes were designed without requiring even spacing along the transcript body, using uniform expression weights and the default filtering criteria for selecting target regions. Probes were generated for all transcripts in the provided MERFISH barcode codebook and multi-FASTA input files.

#### TrueProbes Probe Designer Parameter Settings

TrueProbes probes were designed using probes of 20 nucleotides in length with a minimum spacing of 3 nt between adjacent probes. On- and off-target assessments were restricted to RNA sequences. DNA was excluded, and only BLAST alignments with ≥15 nt of homology were considered. Probe binding affinities were evaluated using the DNA/DNA nearest-neighbor free energy model (46) with 300 mM Na^+^ at 37°C. Probe selection was performed under the equal transcript expression assumption.

#### TrueProbes Designer Thermodynamic-Kinetic Reaction Framework

The TrueProbes platform simulates probe performance using a hybridization reaction model that incorporates key biomolecular interactions affecting smRNA-FISH efficacy (19). The framework considers three primary reaction types: (i) self-dimerization, where a probe binds to itself due to internal self-complementarity; (ii) cross-dimerization, where two distinct probes bind to each other through complementary regions; and (iii) hybridization to target or off-target molecules, where a probe binds a complementary sequence on an intended (on-target) or unintended (off-target) transcript.

#### Probe Alignment Mapping

To evaluate both target hybridization and potential secondary structures, sequence alignments were performed between probes and genome/transcriptome references and between individual probes. On/off-target hybridization was assessed using the NCBI BLAST+ (v2.8.1) algorithm (47) locally in MATLAB. Blast was performed locally with the following parameter settings (E-value threshold 1000, num alignments 1000, word size 7, reward 1, penalty -3, gapopen 5, gapextend 2, and dust low-complexity filtering set to ‘no’). Hits shorter than 15 nucleotides were excluded, as they are unlikely to affect hybridization kinetics significantly. For valid hits, the probe and target sequences, alignment positions, strand orientation, and molecule identifiers were recorded. Self-dimerization and cross-dimerization interactions were evaluated using MATLAB’s oligoprop and swalign functions. Probe binding site maps were constructed based on BLAST alignment data and filtered to ensure non-overlapping binding regions between probes and targets.

#### RNA-seq Data Processing and Alignment for Probe Design Off-Target Quantification

Cell line TPM gene-level and transcript isoform-level expression data from the Broad Institute Cancer Cell Line Encyclopedia (CCLE) Project (38) were analyzed using a methodology adapted from the Human Protein Atlas (41) with protein-coding and non-coding gene TPM trimmed mean of m-value (24) (TMM) normalized across CCLE cell lines to generate normalized transcript per million (nTPM) using MATLAB implementations of NOIseq (49). From gene-level and transcript-level nTPM matrices in MATLAB, cell-line agnostic equal expression levels and average cell-line expression levels were computed as baselines for quantifying probe off-target binding in individual cell lines. Gene expression data indexed using EMBL ENSTx and ENSGx identification were then cross-identified into RefSeq identification numbers used in BLAST to generate software-specific expression matrices containing transcript expression levels for on and off-target RNA for equal, mean average, and cell-line specific. Gene-level expression data was imputed into on/off-target expression levels by normalizing gene-level nTPM by the maximum number of isoforms for each gene across EMBL-EBI and NCBI RefSeq reference genome annotations.

#### Steady State Equilibrium Probe Binding Reactions

All reaction energies were determined using software that computes the Gibbs free energy for any two sequences of oligonucleotides binding to each other at all temperatures within a given temperature range. By considering all the distinct ways a probe can self-dimerize, cross-dimerize, or bind targets, reaction Gibbs free energies were used to compute the dissociation constants for self-hybridization (K_d,Self_), cross-dimerization (K_d,Cross_), and target binding (K_d,Target_) potential probe reactions (33, 34, 45). The dissociation constant for probe (p_i_) self-hybridization 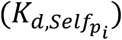 can be seen in Equation 1 using the self-hybridization reaction Gibbs free energy 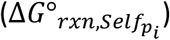 for each potential self-hybridization configuration (L) given the ideal gas constant (R) and the hybridization temperature (T). Each potential self-hybridization configuration is a distinct sequence of sequences within the probe that self-hybridize. The overall dissociation constant for probe (p_i_) self-hybridization 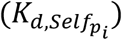 is the sum of the dissociation constants for each potential self-hybridization configuration state L.

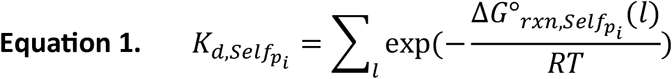

The dissociation constant for probe (p_i_) cross-dimerizing with the probe 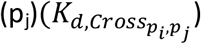 was similarly computed in Equation 2 as the cross-dimerization Gibbs free energy 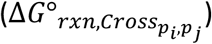 for each potential cross-dimerization configuration state L. Cross-dimerization has two configurations, which are the most energetically favorable potential base-pairs alignment interactions between the two probes are aligning either probe in the paired positions of (5’ to 3’/5’ to 3’) or (5’ to 3’/3’ to 5’). The overall dissociation constant for probe (p_i_) cross-dimerizing with the probe (p_j_) 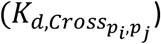 is the sum of the dissociation constants 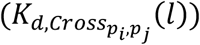 across all potential cross-dimerization configuration states, L.

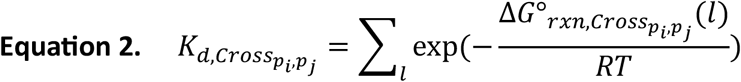

The dissociation constant for probe (p_i_) binding target (J) at the binding site (S) 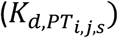 was then computed in Equation 3 using the probe-target site Gibbs free energy 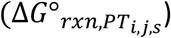 for each potential binding reaction that probe (p_j_) has for the target (J) on the binding site (S). Each probe-target binding site duplex interaction is the paired probe and target sequences from BLAST results within a particular identified binding site (S). The overall dissociation constant for probe (p_i_) binding target (J) at a binding site (S) 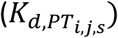 is the sum of the dissociation constants 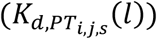 for all interactions (potential probe-target J site S binding configuration state L) the probe might have with that target site.

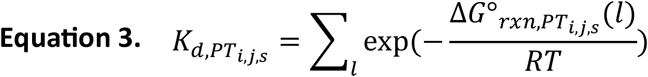

The steady-state concentration of probes bound to targets, cross-dimerized, and self-dimerized, was related to the reaction dissociation constants of probe-target binding 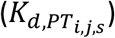, cross-dimerization 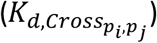 and self-hybridization 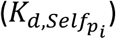 by the mass action equations in Equations 4-6, which describe the steady-state concentration of probes 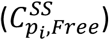 and target sites 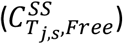 free in-solution to the concentration of self-hybridized 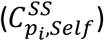, cross-dimerized 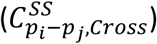, and probes bound to targets 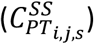.

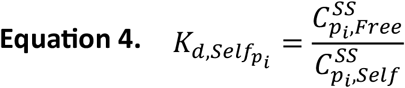

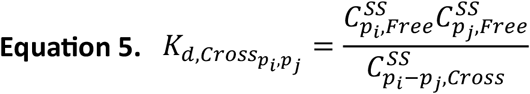

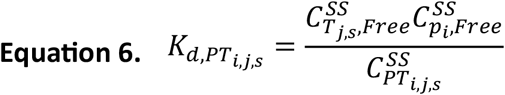

#### TrueProbes smRNA-FISH Probe Selection

Probes to design for smRNA-FISH experiments were prioritized by sequentially sorting probes lowest to highest by their number of off-targets, followed by the estimated difference between on-target binding and the sum of off-target and secondary structure binding affinity, which we denoted as prioritization rank sorting. The default to evaluate probe binding affinity calculations is at 37°C. First, probes with no off targets were selected to design the most probes that cover the target with the highest on-target binding energy corrected for secondary structure binding affinity. For probes without off targets we then performed sequentially the following three-step probe selection process: i) we selected the probe with the highest prioritization rank, ii) all remaining probes that overlap with probes we have chosen already were removed from design consideration, iii) any new probes cross-dimerizing with already selected probes were minimized by updating the prioritization rank of all remaining probes when adding their cross-dimerization affinities to probes already designed to their overall secondary structure binding affinity when evaluating probes prioritization rank. This three-step process was then repeated until no more probes without off-targets could be designed. The remaining probes with off-targets were prioritization rank sorted, and the three-step process was repeated for probes with off-targets based on their number of off-targets, with or without weighting by gene expression levels, until the user-defined maximum number of probes was exceeded or no probes remained to be designed.

#### Steady-State Equilibrium Probe and Target Concentrations

After selecting probes, we performed mass conservation balances on the total concentration of probes and target molecules (50) to compute the equilibrium concentration of probes and targets free in solution, the concentration of probes binding targets, and the probability of any probe binding any target sites. These balances also depend on the parity coefficient defined in Equation 7, which we used to determine the stochiometric concentration of a probe in each cross-dimerization reaction, which we defined as one for reactions between two different probes (i,j), and two for reactions between a probe (i) and another copy of that probe.

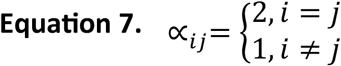

We then generated the initial recursive expression for the steady-state concentration of probes free in-solution 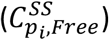 in Equation 8 using the dissociation constants of probe (p_i_) binding target (J) at the binding site (S) 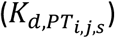 for all target sites it binds, cross-dimerizing dissociation constants 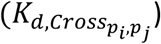 for all probes (p_j_) that probe (p_i_) cross-dimerizes, a self-hybridization constant for probe (p_i_) self-hybridization 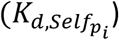, and the total concentration of probe (p_i_)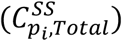 given an initial guess for the steady-state free in-solution concentration of probe (p_i_)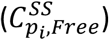, and target (J) site (S) binding sites 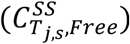 for all target sites probe (p_i_) binds.

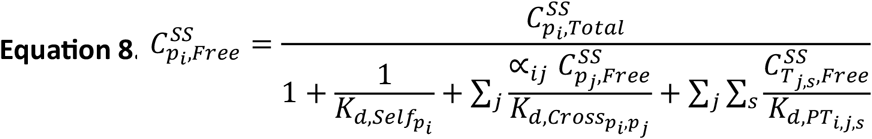

We then determined the free in-solution concentration of RNA target sites explicitly using the concentration of probes free in-solution and target expression levels. The steady-state free concentration of RNA target sites 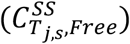 in Equation 9 was derived using the steady-state free in-solution concentration of probes 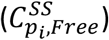, the total expression level of the RNA target (J) site (S)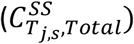, and dissociation constants for all probes (p_i_) binding RNA target (J) site (S)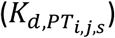. Total expression level of RNA targets was converted to concentration levels in Equation 10 by normalizing nTPM expression values by the volume of a sphere with a radius of 10μm, approximating a Jurkat cell (*R*_*cell*_).

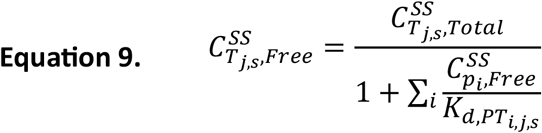

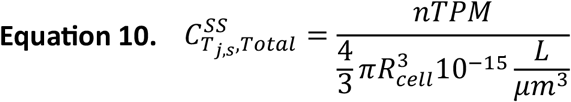

When performing smRNA-FISH probe simulations, we simplified the equation for the probe binding equilibrium to utilize dissociation constants and the concentration of probes free in solution. Combining all the descriptions relating steady-state target (J) site (S) free in-solution concentration 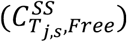 to steady-state free probe in-solution concentration 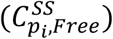, we obtained Equation 11 for the probe’s steady-state free in-solution concentration 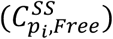 with degrees of freedom (d.o.f) equal to the number of probes designed. We solved this equation in parallel for all probes in a probe set.

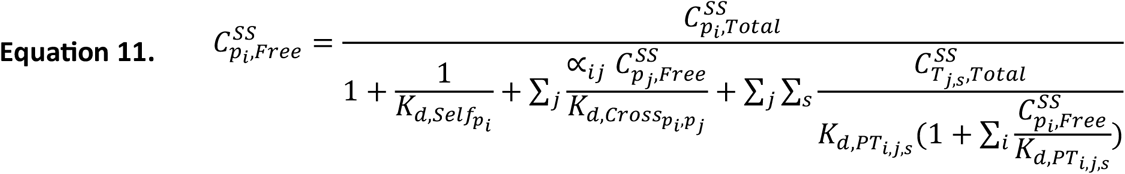

We then vectorized the smRNA-FISH probe-target equilibrium concentration equations to facilitate solving these equilibrium recursive functions given a vector of all probe free-steady state concentrations 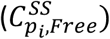 in a probe set (P), which we defined as 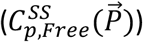 in Equation 12.

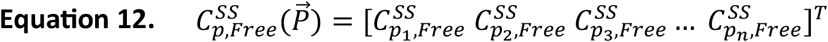

The equation for the concentration of probe free-steady state concentrations 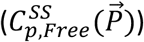was vectorized in Equation 13 using element-wise division, multiplication, and exponentiation with the identity matrix denoted by *I*. For Equation 13, we denoted element-wise division by the Hadamard division symbol (⊘) for two matrices A and B, as each element in A is divided by its corresponding element in 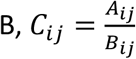, and corresponds to *C* = *A* ⊘ *B* where ⊘ is ./ in MATLAB.

Likewise, we denoted element-wise multiplication by (⊙) for two matrices A and B, as each element in A is multiplied by its corresponding element in B, and is *C* _*ij*_ = *A* _*ij*_ * *B*_*ij*_ and corresponds to *C* = *A* ⊙ *B* where ⊙ is .* in MATLAB. Lastly, we denoted element-wise exponentiation by (∘) for matrices A and some number n, as each element in A is raised to the power of n, where *C*_*ij*_ = *Aij*^*n*^ and corresponds to *C* = *A*^∘*n*^, where ∘ is .^ in MATLAB.

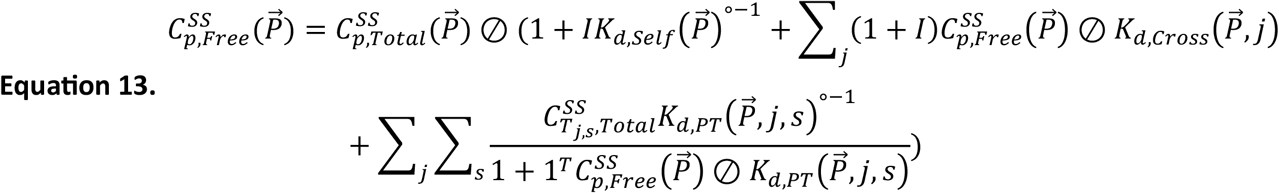

#### Steady-State Equilibrium Probe Reaction Probability

From the steady-state free concentration of probes and targets, we computed the equilibrium concentration of probes binding targets for any number of probes and the probability of probe binding events occurring. Equilibrium equations were rearranged to calculate the probability that probes were self-hybridized, cross-dimerized, or bound to targets. We calculated the probability that a probe binds target (J) at site (S) *P*(*Target*_*js*_ *is Bound*) in Equation 14 by multiplying the steady-state free concentration of target (J) site (S) 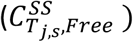 by the total steady-state free concentration of probe (p_i_) 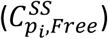 across all probes to generate the overall probability (*P*(*Target*_*js*_ *is Bound*)) that a probe was bound to an individual target (J) binding site (S).

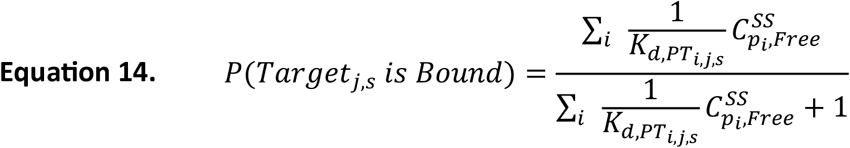

#### Steady-State Probe Target Binding Probability Distributions

We then converted the probability of each site being bound or unbound to compute the probability distribution for the number of probes bound to a target. Because binding sites do not influence each other, the probability distribution for the number of probes bound across all target probe binding sites follows a Poisson-Binomial distribution. The probability distribution for the number of probes bound to a target was derived using a probability-generating function whose characteristic polynomial roots are functions of each binding site probability (51). The Poisson-Binomial generating function in Equation 15 was used to get the relative steady-state probability that each target (J) has any number of probes (n) bound *P*(*n Probes Bound to Target*_*j*_) from the probability that each site was bound *P*(*Target*_*js*_ *is Bound*) and the total number of target (J) probe binding sites N_sites_(J).

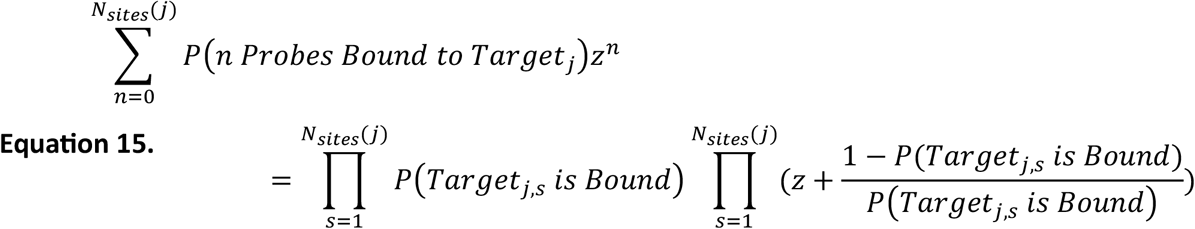

We then determined the probability that on-/off-targets have any number of probes (n) bound in different reference cell backgrounds by bringing reference expression levels of individual genes (J) back into the calculation. We then solved, with or without gene expression levels, for the number of on-targets with n probes bound (*N*_*on*_(*n*)) and the number of off-targets with n probes bound (*N*_*off*_(*n*)) in Equations 16 and 17, respectively, which combined the expression level of each target (*Expression*_*j*_) with the probability that each on or off-target (J) has n probes bound *P*(*n Probes Bound to Target*_*j*_). Following this, we translated the number of off-targets with n bounds (*N*_*off*_(*n*)) bound into a probe set specificity load in Equation 18, describing the number of probes bound to off-target transcripts compared to the mean number of probes bound to the on-target.

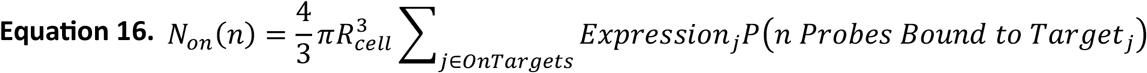

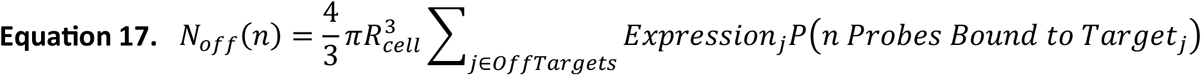

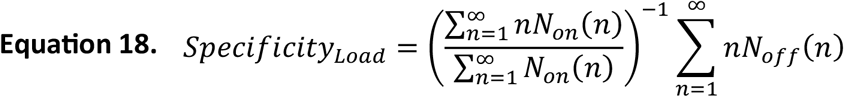

#### smRNA-FISH Cell Segmentation and Spot Detection

smRNA-FISH spots were detected using TrueSpot, an automated smRNA-FISH spot detection software (52, doi: https://doi.org/10.1101/2025.01.10.632467). Before spots were detected, an updated command line version of CellDissect (53) in TrueSpot was used for cell segmentation in all smRNA-FISH images. CellDissect segments nuclei in 3D and cells in 2D using the TRANS and DAPI channel images. CellDissect uses the DAPI signal of the nucleus to generate automated thresholds to quantify individual non-overlapping nuclei used in determining cell boundaries.

After cell segmentation, for each smRNA-FISH CY5 channel image, the pipeline performed dead pixel removal to reduce false positives, followed by a Laplacian of Gaussian (LoG) transformation applied to each slice of the raw 3D image stack and edge detection. TrueSpot then used a range of intensity thresholds to find local intensity maxima, i.e., potential spots, within each filtered image stack at each intensity threshold. For each smRNA-FISH image, TrueSpot constructed a spot detection curve by counting the number of spots detected at each threshold. These curves were log10-transformed and analyzed using two-piece linear regression and the median absolute deviation (MAD) to estimate optimal thresholds and their ranges. A final detection threshold was selected using a weighted combination of the linear regression fit and MAD-based metrics. TrueSpot was run under two settings in our analyses: (i) Image-specific thresholding, where a separate detection threshold was calculated for each 3D image stack. (ii) Global thresholding, where all images used the minimum threshold derived from the individual image analyses.

#### smRNA-FISH Spot Quantification

smRNA-FISH spot and background intensities were quantified using TrueSpot at both the single-cell and single-spot levels. Spot quantification was performed using the minimum image-level threshold across all smFISH images. For each detected spot, TrueSpot fitted a Gaussian to the surrounding voxels around each spot slice-by-slice in the original RNA-FISH image stack, then integrated the signal across z-stacks to generate a 3D punctum. Spot intensity was calculated as the total integrated intensity of all voxels within the 3D region.

TrueSpot then computed the centroid of each spot and applied 2D Gaussian fits across ±2 z-stacks around the centroid to determine spot amplitude, fitted intensity, radius, and a spot mask. Before intensity quantification, all images were corrected for uneven illumination using a reference correction matrix. This matrix was generated by 2D smoothing of the percent maximum pixel intensity at each position across z-stacks in cell-free control images.

For each spot, TrueSpot calculated a set of intensity statistics: minimum, maximum, mean, median, standard deviation, MAD, and intensity histograms. Background was measured locally for each detected spot by computing the mean, median, and maximum intensity of all pixels not associated with any detected spot within ± 5 pixels in the x/y-directions from the spots Gaussian fit x/y full width at half maximum across ±2 z-stack around the centroid of the spot. While both maximum and mean spot intensities were recorded, maximum intensity was primarily used for quantification. TrueSpots then merged nearby local maxima detections—spots within one pixel in the x, y, or z direction—into actual and duplicate candidate smRNA-FISH spots.

For all RNA spot intensity quantifications, software and replica specific spot detection thresholds were used based on TrueSpot automated thresholding. For each software RNA-FISH image, TrueSpot generated eighteen threshold suggestions. For each software in each replica, these suggestions were then aggregated, and their median was applied to detect spots across all fields of view images at each replica. All spots detected at these image level thresholds were then filtered to focus only on non-duplicate spots.

From these values, TrueSpot generated distributions of maximum spot intensities and mean background intensities. Signal-minus-background was calculated as the difference between the two. Finally, the signal-to-noise ratio (SNR) was computed for each spot using Equation 19, where μ_signal_ is the maximum spot intensity, μ_background_ is the mean background intensity, and σ_background_ is the background standard deviation. This per-pixel SNR accounts for both the absolute signal level and variability in background, providing a robust measure of contrast in fluorescent microscopy images.

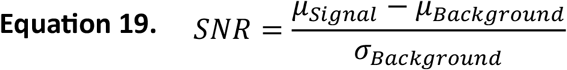

After quantifying spot intensity metrics, the distribution of total smRNA-FISH spot counts per cell was computed as detected by TrueSpot for each probe design software. Given the generally low number of spots per cell, our analysis was focused on total spot counts per cell rather than per-spot metrics. To minimize artifacts, several quality filters were applied. Cells with abnormally large volumes—typically resulting from poor segmentation—were excluded. Lysed or damaged cells exhibiting unusually high mean background intensity (greater than 1000 a.u. per pixel) and artifactual spots with radii smaller than half a pixel were removed. The total number of smRNA-FISH spots for each remaining cell was calculated by counting all spots whose dropout threshold met or exceeded the cell’s spot detection threshold.

#### Statistics and Reproducibility

RNA-FISH experiments were conducted with three separate replicas, each with at least three fields of view per software per replica. All intensity and spot count distributions were analyzed using two-sample Kolmogorov-Smirnov tests, Kruskal-Wallis tests and Analysis of Variance (ANOVA). Multiple comparison tests were performed with MATLAB function multcompare with an Alpha value of 0.05 using Bonferroni critical value correction to compare metric distributions for pooled replica data in Figure 4 and individual replica data in Figure S5. All hypothesis testing ANOVA tables, multiple comparison tests, with their associated P-values and group mean estimates and standard errors are in Supplemental Excel Spreadsheet Table 1 for individual replica, replica pooled, and pairwise software and replica data comparisons.

## Resource Availability

### Lead contact

All requests for further information on either the TrueProbes smRNA-FISH design software or the smRNA-FISH experiments should be directed to the lead contact, Gregor Neuert

## Data and Code Availability

All pre-processed raw images can be accessed at the BioImage Archive (https//www.ebi.ac.uk/biostudies/studies/). All MATLAB-processed experimental image data and TrueProbes probe design computational model prediction data can be accessed at <u>Zenodo</u> (https://zenodo.org/records/). All RNA-FISH software, *ARF4* probe set, experimental statistical comparisons, and probe sequences are included in the supplemental Excel spreadsheet tables 1 and 2, respectively. TrueProbes smRNA-FISH probe design software and smRNA-FISH probe set prediction software, with documentation and sample output, are available at GitHub (https://github.com/neuertlab/TrueProbes). All sample data, gene transfer format (GTF) and generic feature format (GFF) annotation files, reference genome, transcriptome, and gene expression data currently used in TrueProbes are available at GitHub (https://github.com/neuertlab/TrueProbes_Data). The TrueSpot software used for spot detection is available at GitHub (https://github.com/neuertlab/TrueSpot) and <u>Zenodo</u> (https://zenodo.org/records/13345321).

## Author Contributions

Conceptualization, G.N., B.K.; methodology, G.N., B.K., J.H.; software, G.N., B.K., J.H., B.H.; resources, J.A.; formal analysis, J.H., B.H.; investigation, J.H., J.A.; data curation, J.A., B.K., B.H.; writing – original draft, J.H., writing – review & editing, G.N., J.H.; validation, J.A.; visualization, G.N., J.H.; supervision, G.N.; project administration, G.N.; funding acquisition, G.N.

## Acknowledgements

We want to thank Dr. Steven J. Walker and Dr. Antentor O. Hinton for their feedback on the manuscript writing. We would additionally like to thank Dr. Manuel Ascano and Samantha Lisy for their help with HPLC purification of RNA-FISH probes.

## References

1. Pang, B., van Weerd, J. H., Hamoen, F. L., and Snyder, M. P. (2023) Identification of non-coding silencer elements and their regulation of gene expression Nat Rev Mol Cell Biol 24, 383–395

2. Unger Avila, P., Padvitski, T., Leote, A. C., Chen, H., Saez-Rodriguez, J., Kann, M. et al. (2024) Gene regulatory networks in disease and ageing Nat Rev Nephrol 20, 616–633

3. Alseekh, S., Aharoni, A., Brotman, Y., Contrepois, K., D’Auria, J., Ewald, J. et al. (2021) Mass spectrometry-based metabolomics: a guide for annotation, quantification and best reporting practices Nat Methods 18, 747–756

4. Satam, H., Joshi, K., Mangrolia, U., Waghoo, S., Zaidi, G., Rawool, S. et al. (2023) Next-Generation Sequencing Technology: Current Trends and Advancements Biology (Basel) 12,

5. Elowitz, M. B., Levine, A. J., Siggia, E. D., and Swain, P. S. (2002) Stochastic gene expression in a single cell Science 297, 1183–1186

6. Ozbudak, E. M., Thattai, M., Kurtser, I., Grossman, A. D., and van Oudenaarden, A. (2002) Regulation of noise in the expression of a single gene Nature Genetics 31, 69–73

7. Desai, R. V., Chen, X., Martin, B., Chaturvedi, S., Hwang, D. W., Li, W. et al. (2021) A DNA repair pathway can regulate transcriptional noise to promote cell fate transitions Science 373,

8. Neuert, G., Munsky, B., Tan, R. Z., Teytelman, L., Khammash, M., and van Oudenaarden, A. (2013) Systematic Identification of Signal-Activated Stochastic Gene Regulation Science 339, 584–587

9. Vandereyken, K., Sifrim, A., Thienpont, B., and Voet, T. (2023) Methods and applications for single-cell and spatial multi-omics Nature Reviews Genetics

10. Heumos, L., Schaar, A. C., Lance, C., Litinetskaya, A., Drost, F., Zappia, L. et al. (2023) Best practices for single-cell analysis across modalities Nature Reviews Genetics 24, 550–572

11. Khetan, N., Zuckerman, B., Calia, G. P., Chen, X., Garcia Arceo, X., and Weinberger, L. S. (2024) Single-cell RNA sequencing algorithms underestimate changes in transcriptional noise compared to single-molecule RNA imaging Cell Rep Methods 4, 100933

12. Braselmann, E., Department of Biochemistry, B. I., University of Colorado Boulder, 3415 Colorado Avenue, Boulder, CO 80309, USA, Rathbun, C., Department of Biochemistry, B. I., University of Colorado Boulder, 3415 Colorado Avenue, Boulder, CO 80309, USA, Richards, E. M., Department of Biochemistry, B. I., University of Colorado Boulder, 3415 Colorado Avenue, Boulder, CO 80309, USA et al. (2020) Illuminating RNA Biology: Tools for Imaging RNA in Live Mammalian Cells Cell Chemical Biology 27, 891–903

13. Femino, A. M., Fay, F. S., Fogarty, K., and Singer, R. H. (1998) Visualization of single RNA transcripts in situ Science 280, 585–590

14. Raj, A., van den Bogaard, P., Rifkin, S. A., van Oudenaarden, A., and Tyagi, S. (2008) Imaging individual mRNA molecules using multiple singly labeled probes Nature Methods 5, 877–879

15. Chen, K. H., Boettiger, A. N., Moffitt, J. R., Wang, S., and Zhuang, X. (2015) RNA imaging. Spatially resolved, highly multiplexed RNA profiling in single cells Science 348, aaa6090

16. Shah, S., Lubeck, E., Zhou, W., and Cai, L. (2016) In Situ Transcription Profiling of Single Cells Reveals Spatial Organization of Cells in the Mouse Hippocampus Neuron 92, 342–357

17. Tsanov, N., Samacoits, A., Chouaib, R., Traboulsi, A.-M., Gostan, T., Weber, C. et al. (2016) smiFISH and FISH-quant–a flexible single RNA detection approach with super-resolution capability Nucleic acids research 44, e165–e165

18. Moffitt, J. R., Lundberg, E., and Heyn, H. (2022) The emerging landscape of spatial profiling technologies Nature Reviews Genetics 23, 741–759

19. Liu, G., and Zhang, T. (2021) Single copy oligonucleotide fluorescence in situ hybridization probe design platforms: development, application and evaluation International Journal of Molecular Sciences 22, 7124

20. Coassin, S. R., Orjalo, A. V., Semaan, S. J., and Johansson, H. E. (2014) Simultaneous detection of nuclear and cytoplasmic RNA variants utilizing Stellaris® RNA fluorescence in situ hybridization in adherent cells In Situ Hybridization Protocols 189–199

21. Safieddine, A., Coleno, E., Lionneton, F., Traboulsi, A. M., Salloum, S., Lecellier, C. H. et al. (2023) HT-smFISH: a cost-effective and flexible workflow for high-throughput single-molecule RNA imaging Nature Protocols 18, 157–187

22. Hershberg, E. A., Camplisson, C. K., Close, J. L., Attar, S., Chern, R., Liu, Y. et al. (2021) PaintSHOP enables the interactive design of transcriptome- and genome-scale oligonucleotide FISH experiments Nature Methods 2021 18:8 18, 937–944

23. Kim, S. W., Hayashi, M., Lo, J. F., Yang, Y., Yoo, J. S., and Lee, J. D. (2003) ADP-ribosylation factor 4 small GTPase mediates epidermal growth factor receptor-dependent phospholipase D2 activation J Biol Chem 278, 2661–2668

24. Robinson, M. D., and Oshlack, A. (2010) A scaling normalization method for differential expression analysis of RNA-seq data Genome Biol 11, R25

25. Arvey, A., Hermann, A., Hsia, C. C., Ie, E., Freund, Y., and McGinnis, W. (2010) Minimizing off-target signals in RNA fluorescent in situ hybridization Nucleic Acids Res 38, e115

26. Beliveau, B. J., Kishi, J. Y., Nir, G., Sasaki, H. M., Saka, S. K., Nguyen, S. C. et al. (2018) OligoMiner provides a rapid, flexible environment for the design of genome-scale oligonucleotide in situ hybridization probes Proceedings of the National Academy of Sciences 115, E2183–E2192

27. Andersson, P., Burel, S. A., Estrella, H., Foy, J., Hagedorn, P. H., Harper, T. A. et al. (2025) Assessing Hybridization-Dependent Off-Target Risk for Therapeutic Oligonucleotides: Updated Industry Recommendations Nucleic Acid Ther 35, 16–33

28. S, S., S, S., and AC, P. (2022) Detecting RNA-RNA interactome Wiley interdisciplinary reviews RNA 3,

29. Lima, J. F., Maia, P.T., Magalhães, B., Cerqueira, L., and Azevedo, N. F. (2020) A comprehensive model for the diffusion and hybridization processes of nucleic acid probes in fluorescence in situ hybridization Biotechnology and Bioengineering 117, 3212–3223

30. Todisco, M., and Szostak, J. (2022) Hybridization kinetics of out-of-equilibrium mixtures of short RNA oligonucleotides Nucleic Acids Research 50, 9647–9662

31. Marimuthu, K., and Chakrabarti, R. (2014) Sequence-dependent theory of oligonucleotide hybridization kinetics J Chem Phys 140, 175104

32. Wright, E. S., Yilmaz, L. S., Corcoran, A. M., Ökten, H. E., and Noguera, D. R. (2014) Automated design of probes for rRNA-targeted fluorescence in situ hybridization reveals the advantages of using dual probes for accurate identification Appl Environ Microbiol 80, 5124–5133

33. Oliveira, L. M., Long, A. S., Brown, T., Fox, K. R., and Weber, G. (2020) Melting temperature measurement and mesoscopic evaluation of single, double and triple DNA mismatches Chemical Science 11, 8273–8287

34. de Oliveira Martins, E., and Weber, G. (2024) Nearest-neighbour parametrization of DNA single, double and triple mismatches at low sodium concentration Biophys Chem 306, 107156

35. Goldfarb, T., Kodali, V. K., Pujar, S., Brover, V., Robbertse, B., Farrell, C. M. et al. (2025) NCBI RefSeq: reference sequence standards through 25 years of curation and annotation Nucleic Acids Res 53, D243–D257

36. Dyer, S. C., Austine-Orimoloye, O., Azov, A. G., Barba, M., Barnes, I., Barrera-Enriquez, V. P. et al. (2025) Ensembl 2025 Nucleic Acids Res 53, D948–D957

37. Mudge, J. M., Carbonell-Sala, S., Diekhans, M., Martinez, J. G., Hunt, T., Jungreis, I. et al. (2025) GENCODE 2025: reference gene annotation for human and mouse Nucleic Acids Res 53, D966–D975

38. Ghandi, M., Huang, F. W., Jané-Valbuena, J., Kryukov, G. V., Lo, C. C., McDonald, E. R. et al. (2019) Next-generation characterization of the Cancer Cell Line Encyclopedia Nature 569, 503–508

39. Consortium, G. (2020) The GTEx Consortium atlas of genetic regulatory effects across human tissues Science 369, 1318–1330

40. Tomczak, K., Czerwińska, P., and Wiznerowicz, M. (2015) The Cancer Genome Atlas (TCGA): an immeasurable source of knowledge Contemp Oncol (Pozn) 19, A68–77

41. Jin, H., Zhang, C., Zwahlen, M., von Feilitzen, K., Karlsson, M., Shi, M. et al. (2023) Systematic transcriptional analysis of human cell lines for gene expression landscape and tumor representation Nat Commun 14, 5417

42. Consortium, T. M. (2018) Single-cell transcriptomics of 20 mouse organs creates a Tabula Muris Nature 562, 367–372

43. Jones, R. C., Karkanias, J., Krasnow, M. A., Pisco, A. O., Quake, S. R., Salzman, J. et al. (2022) The Tabula Sapiens: A multiple-organ, single-cell transcriptomic atlas of humans Science 376, eabl4896

44. Allawi, H. T., and SantaLucia, J. (1997) Thermodynamics and NMR of internal G.T mismatches in DNA Biochemistry 36, 10581–10594

45. SantaLucia Jr, J., and Hicks, D. (2004) The thermodynamics of DNA structural motifs Annu Rev Biophys Biomol Struct 33, 415–440

46. Sugimoto, N., Nakano, S., Yoneyama, M., and Honda, K. (1996) Improved thermodynamic parameters and helix initiation factor to predict stability of DNA duplexes Nucleic Acids Res 24, 4501–4505

47. Boratyn, G. M., Camacho, C., Cooper, P. S., Coulouris, G., Fong, A., Ma, N. et al. (2013) BLAST: a more efficient report with usability improvements Nucleic acids research 41, W29–W33

48. Cock, P. J., Antao, T., Chang, J. T., Chapman, B. A., Cox, C. J., Dalke, A. et al. (2009) Biopython: freely available Python tools for computational molecular biology and bioinformatics Bioinformatics 25, 1422–1423

49. Tarazona, S., Furió-Tarí, P., Turrà, D., Pietro, A. D., Nueda, M. J., Ferrer, A. et al. (2015) Data quality aware analysis of differential expression in RNA-seq with NOISeq R/Bioc package Nucleic Acids Res 43, e140

50. Horne, M. T., Fish, D. J., and Benight, A. S. (2006) Statistical thermodynamics and kinetics of DNA multiplex hybridization reactions Biophysical Journal 91, 4133–4153

51. Fernández, M., and Williams, S. (2010) Closed-form expression for the poisson-binomial probability density function IEEE Transactions on Aerospace and Electronic Systems 46, 803–817

52. Hospelhorn, B. G., Kesler, B. K., Jashnsaz, H., and Neuert, G. (2025) TrueSpot: A robust automated tool for quantifying signal puncta in fluorescent imaging bioRxiv

53. Kesler, B. K., Adams, J., and Neuert, G. (2025) Transcriptional stochasticity reveals multiple mechanisms of long non-coding RNA regulation at the Xist-Tsix locus Nat Commun 16, 4223

